# Jumping between Turtles, Fishes, and a Frog: The Unexpected Horizontal Transfer of a DNA Transposon

**DOI:** 10.1101/2023.08.19.550906

**Authors:** Nozhat T. Hassan, James D. Galbraith, David L. Adelson

## Abstract

Horizontal transfer of transposable elements (HTT) has been reported across many species and the impact of such events on genome structure and function has been well described. However, few studies have focused on reptilian genomes, especially HTT events in Testudines (turtles). Here, we investigated the repetitive content of *Malaclemys terrapin terrapin* (Diamondback turtle) and found a high similarity hAT-6 DNA transposon shared between other turtle species, ray-finned fishes, and a frog. hAT-6 was notably absent in taxa closely related to turtles, such as crocodiles and birds. Successful invasion of DNA transposons into new genomes requires the conservation of specific residues in the encoded transposase, and through structural analysis, these residues were identified indicating retention of functional transposition activity. We document a rare and recent HTT event of a DNA transposon between turtles which are known to have a low genomic evolutionary rate and ancient repeats.

## Introduction

A large proportion of eukaryotic genomes is composed of mobile repetitive sequences known as transposable elements (TEs). TEs are divided into two classes based on their method of mobilisation. Retrotransposons (Class I) copy themselves using an RNA intermediate which is reverse-transcribed into cDNA and is integrated back into the host genome, whereas DNA transposons (Class II) move via a cut-and-paste mechanism mediated by an encoded transposase [1,2]. One of the largest and most widespread superfamilies of DNA transposons are hAT elements. While there is sequence variation between hAT transposons, they are defined by ∼8 bp terminal target site duplication (TSDs) and ∼15 bp terminal inverted repeats (TIRs) [3]. The hAT transposase is a multidomain protein and the only hAT transposase crystal structure is from the Hermes transposase of the house fly (*Musca domestica*), which showed the presence of conserved residues and domains required for TIR recognition and transposition [4,5].

Testudines (turtles) are a group of reptiles found in diverse ecological settings ranging from terrestrial, marine, and freshwater environments and are a sister group to Archosauria (birds and crocodilians) that diverged during the Permian-Triassic period approximately 257.4 Mya [6,7]. The repetitive content of turtle genomes has not been studied extensively, but current studies conclude that TEs make up approximately 30% of the genome in various turtles and are predominantly LINEs and DNA transposons [8–10]. Turtle genomes generally do not show significant variation in size and appear to evolve slowly compared to other reptiles which makes analysis of repeats an area of interest [11]. Until now, Horizontal transfer of transposable elements (HTT) has not been documented between turtles.

Horizontal transfer is the process by which genetic material is obtained from non-parental individuals, as opposed to vertical transfer which is from parent to offspring [12,13]. TEs, in particular, are widely spread through horizontal transfer in eukaryotes and can persist within the invaded genome [14]. HTT has been documented in several species, for example, SPIN (DNA TEs) and BovB elements (non-LTR retrotransposons) have colonised many squamate reptiles [15–17]. However, in both studies, evidence for HTT was notably absent in turtles. In 2020, a study demonstrated HTT of both classes of TEs between turtles, fishes, and lizards, however, they did not specifically detect hAT-6 HTT events between species or a HTT event between two turtles, or carry out an evolutionary analysis (Zhang et al. 2020). Here in this study, we outline a rare horizontal transfer of a hAT-6 DNA TE between two turtles, a frog, and fishes.

## Results and Discussion

### HTT amongst species of turtles, fishes, and a frog

Horizontal transposon transfer has been widely reported in the species discussed in this study, with new reports emerging describing HTT of both DNA and RNA transposable elements in ray-finned fish, amphibians, and reptiles [18,19]. HTT between turtles, especially of DNA transposons, has been proposed to be a rare event based on our current understanding of turtle genome evolution [18-20]. We have identified the first case of horizontal transfer of a hAT-6 DNA transposon into turtles, ray-finned fishes, and a frog.

We show that hAT-6 TEs are distinct from other hAT and DNA TEs (fig. 1). In addition, the high sequence similarity of TEs from distant species, absence in closely related species, and discordant topologies of the species and hAT-6 phylogenies (fig. 2) make a compelling case for the horizontal transfer of a DNA TE. The first detected transfer of hAT-6 was between *Xenopus tropicalis* (Western clawed frog) and *M. t. terrapin* as it satisfied the previous criteria for HTT. A further 7 hAT-6s were found across turtles, frogs, and fish. In addition, given the high sequence identity between pairs of species (fig. 1), hAT-6 may have repeatedly transferred across aquatic animals over time. While we could not determine the direction of transfer, it is likely to be mediated by a parasitic donor.

**Figure 1:**
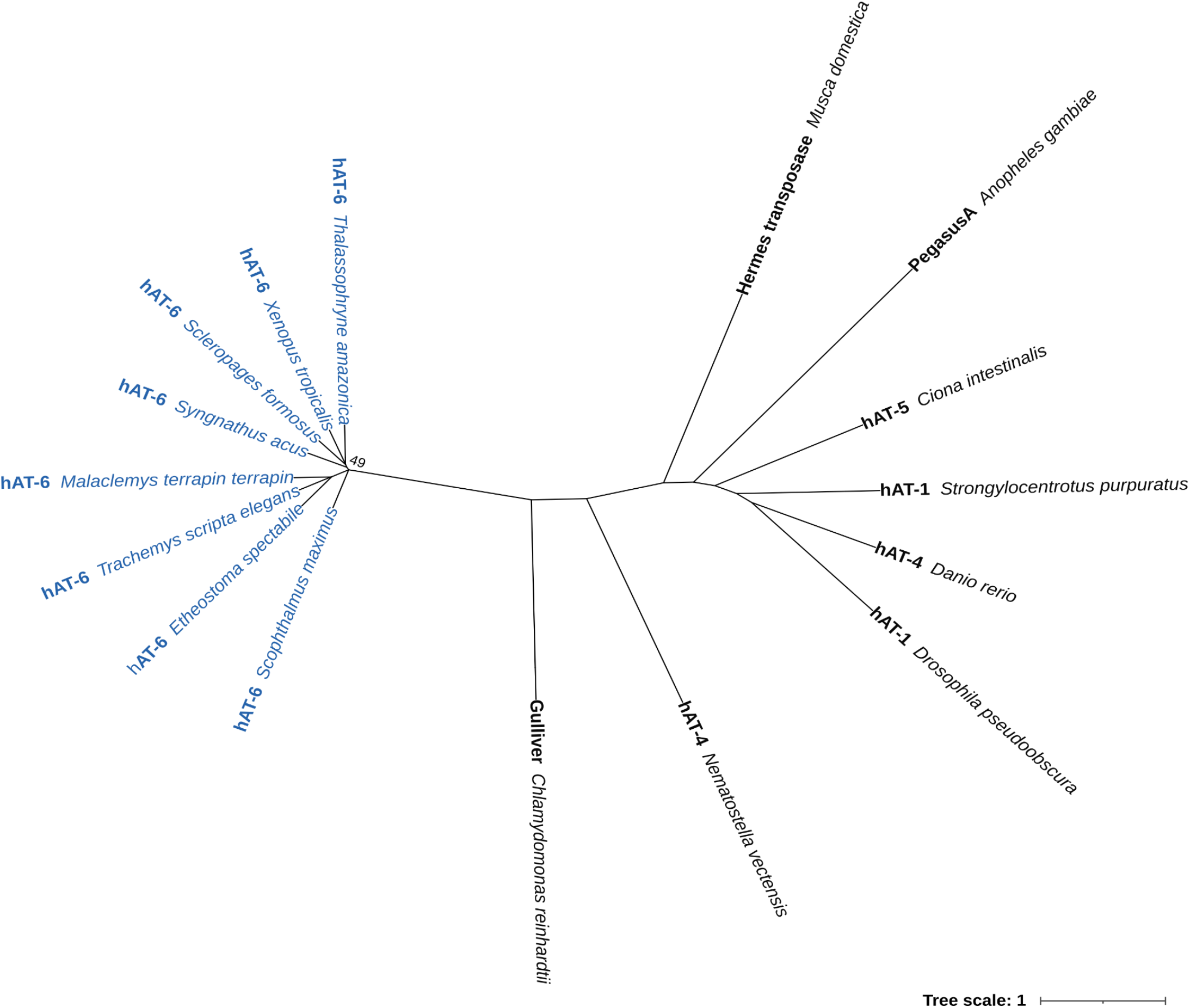
Phylogeny of horizontally transferred hAT-6 TEs in blue text with a sample of other hAT TEs from Repbase. Tree constructed using IQTree (1000 bootstraps) based on MAFFT protein alignments trimmed using Clipkit and plotted in iTOL. Support values under 70 are displayed at nodes.

**Figure 2:**
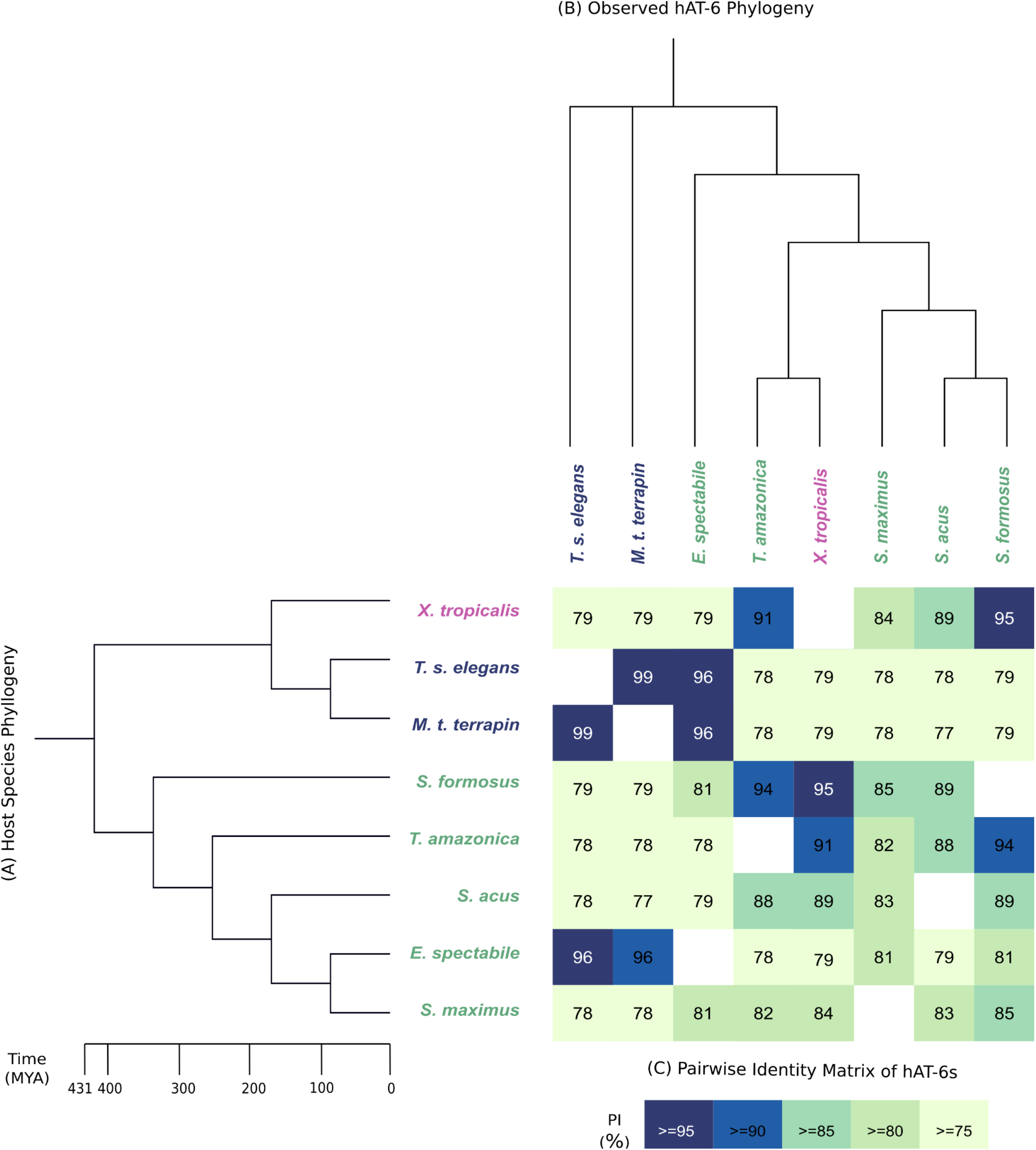
Sequence homology and phylogeny of hAT-6 relative to host species phylogeny. (A) Host species phylogeny of hAT-6 host species. The time of divergence between host species is shown in MYA. (B) Observed hAT-6 phylogeny constructed using IQTree (1000 bootstraps) based on MAFFT protein alignments trimmed using Clipkit and plotted in iTOL. (C) BLASTN pairwise identity matrix of hAT-6 sequence alignments. Darker shading indicates a higher pairwise identity between two hAT-6s. Frogs are in pink text, turtles are in blue text, and fishes are in green text.

**Figure 3:**
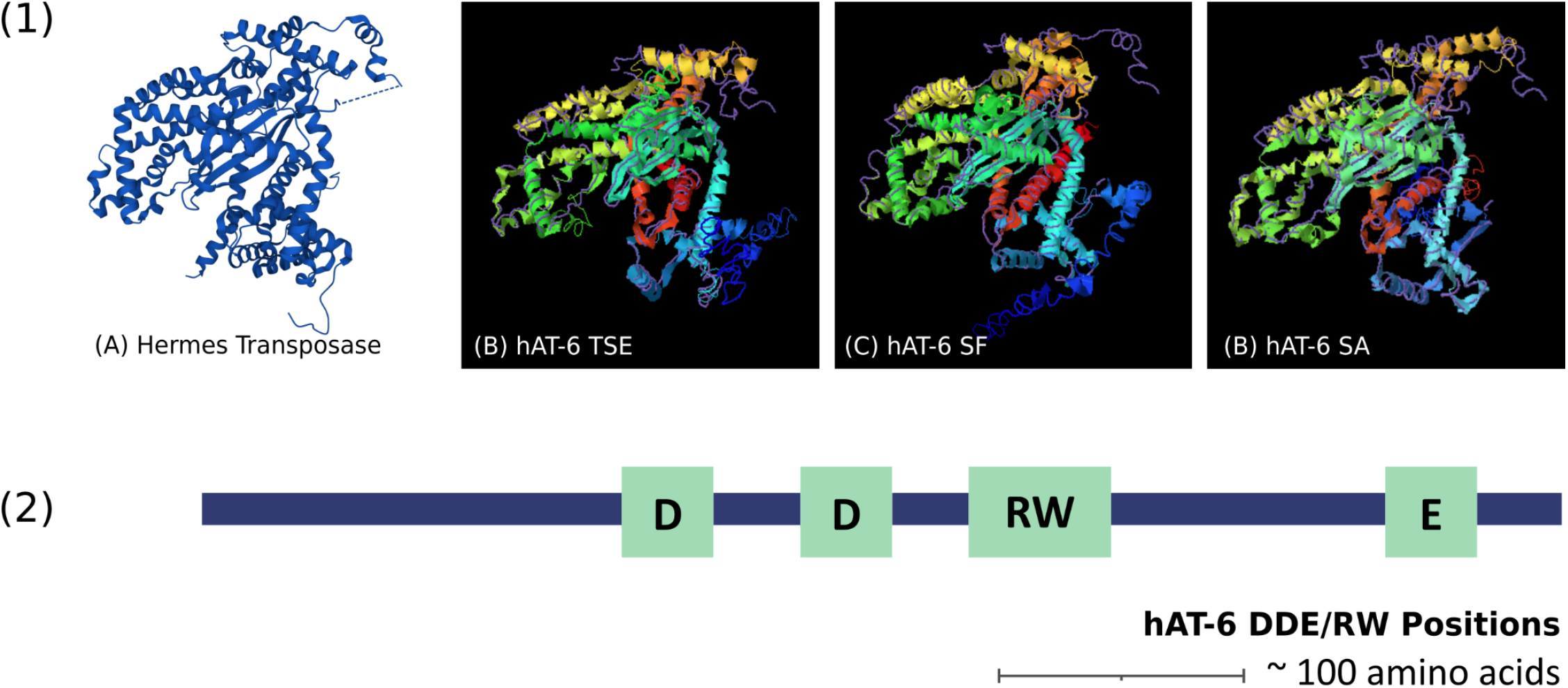
Predicted 3D structure of hAT-6s and the essential residues required for transposition. (1) The predicted 3D structures of a sample of hAT-6 transposases. A) Three-dimensional ribbon structure of the Hermes DNA transposase monomer (https://www.rcsb.org/structure/2BW3). I-TASSER structural alignment of the best-predicted ribbon structures of B) hAT-6 TSE C) hAT-6 SF, and D) hAT-6 SA is overlaid with the Hermes DNA transposase (purple wire), showing high structural similarity. (2) MUSCLE multiple alignment of DDE/RW residue containing regions of autonomous hAT-6 ORFs. DDE/RW residues are highlighted in blue.

### The structure of hAT-6 transposases indicates activity

To support our case for HTT of hAT-6, we determined the predicted structure and the presence of functional domains required for mobility in the corresponding encoded transposases. The conservation of the TIRs/TSDs in full-length hAT-6 transposons is expected and required for HTT. In contrast, the absence of those features in fragmented hAT-6 transposons indicates degradation in the genome. We find conservation of such features in all hAT-6s found (supplementary materials). In addition, the functional annotation of hAT-6 transposases shows the presence of the essential DDE catalytic triad required for transposition and for HTT (fig. 4) [4]. These findings suggest that the hAT-6s have the necessary sequence features and motifs to be horizontally transferred.

**Figure 4:**
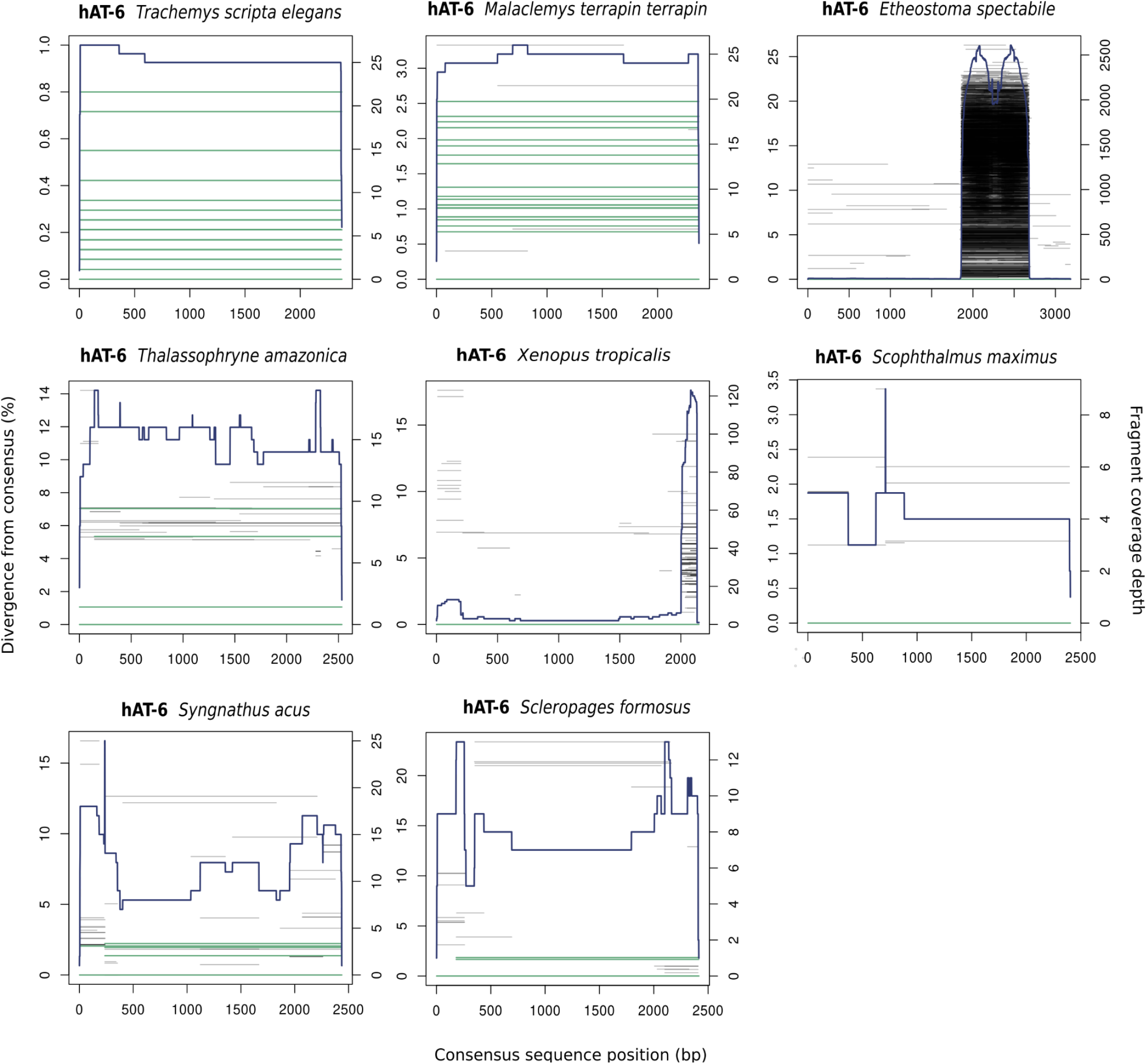
Coverage and divergence plots of the 8 horizontally transferred hAT-6 transposons. hAT-6 relative divergence and abundance were plotted using TE Aid (https://github.com/clemgoub/TE-Aid) [44]. The blue line represents the depth of coverage (right-hand Y-axis) of each fragment aligned to the repeat consensus sequence. Green lines represent a full-length copy of the repeat. Black lines show repeat fragments. Percentage divergence from the consensus sequence is shown on the left-hand Y-axis.

The high percentage identity and low divergence of hAT-6 between species are likely the result of independent transfers from an unknown donor(s). The species in this study are largely found in semi-aquatic or aquatic environments but for the most part, are geographically distant. This indicates possible donor(s) that is/are likely ubiquitously aquatic and/or parasitic in nature. This finding also supports the frequent and recent transfer of transposons in aquatic environments [18,19]. Horizontal transfer is known to be facilitated by parasite-host relationships with recent findings showing HTT of a retrotransposon from parasitic nematodes [21]. Pathogens affecting fish, frogs and turtles include nematodes, tapeworms, flukes, trematodes, and myxosporean parasites [22–24]. The myxozoan endoparasite of the common carp (*Thelohanellus kitaueI*) contained a hAT DNA transposon with motifs remarkably similar to the horizontally transferred hAT-6 TSDs, TIRs, and identical DDE/RW residues. Although there is insufficient evidence to explain the HTT of hAT-6 via a parasitic intermediate, *T. kitauel* shows potential as a parasitic donor as it is known to infect fish and possibly frogs and turtles. As there is geographical overlap between the turtles and *Etheostoma spectabile* (Orangethroat darter, North America), but not between *X. tropicalis* and the other fishes, more genomic data from geographically/ecologically overlapping species are required to determine possible donors (Supplementary materials).

### hAT-6 expansion and divergence

To understand the evolution of each hAT-6 transfer in the host species, we investigated the coverage of each element. The genome coverage plots of hAT-6 in turtles (fig. 4) show very low divergence which may be a product of the slow genome evolution of turtles rather than recent HTT events [19]. But when combined with how unexpectedly similar hAT-6s are across all species, there is evidence pointing towards HTT in turtles. In comparison to the other species, turtles also contained more full-length copies of hAT-6. It is possible that the horizontal transfer of hAT-6 occurred prior to the divergence of *Trachemys scripta elegans* (red-eared slider) and *M. t. terrapin* ∼14MYA, and has been well preserved in the genomes of turtles (*T. s. elegans* and *M. t. terrapin*) because of their low neutral substitution rates [7,19,25]. Without data from closely related taxa, we cannot resolve whether HTT of hAT-6 occurred independently into each turtle species or into their common ancestor.

In *E. spectabile*, where hAT-6 divergence is much greater, we also observed a highly amplified region of ∼800 bp from hAT-6 (fig. 5). Upon further investigation, we found that this region is a hAT-6 derived Miniature Inverted-repeat Transposable Element (MITE). By looking at the Kimura-based divergence of both the hAT-6 and hAT-6 derived MITE from *E. spectabile*, we see that copy number increase of the MITE occurred at the same time as copy number increase of hAT-6 (fig. 5). We also observed two additional instances of hAT-6 copy number increase in E. *spectabile*, however, this was also the case in all instances where hAT-6 was present, which could indicate reactivation of hAT-6 rather than recent HTT (Supplementary materials). However, taken together with other evidence of HTT, we hypothesise that there has been repeated HTT into *E. spectabile*, rather than the more diverged copies of hAT-6 being a degraded version of a similar hAT.

**Figure 5:**
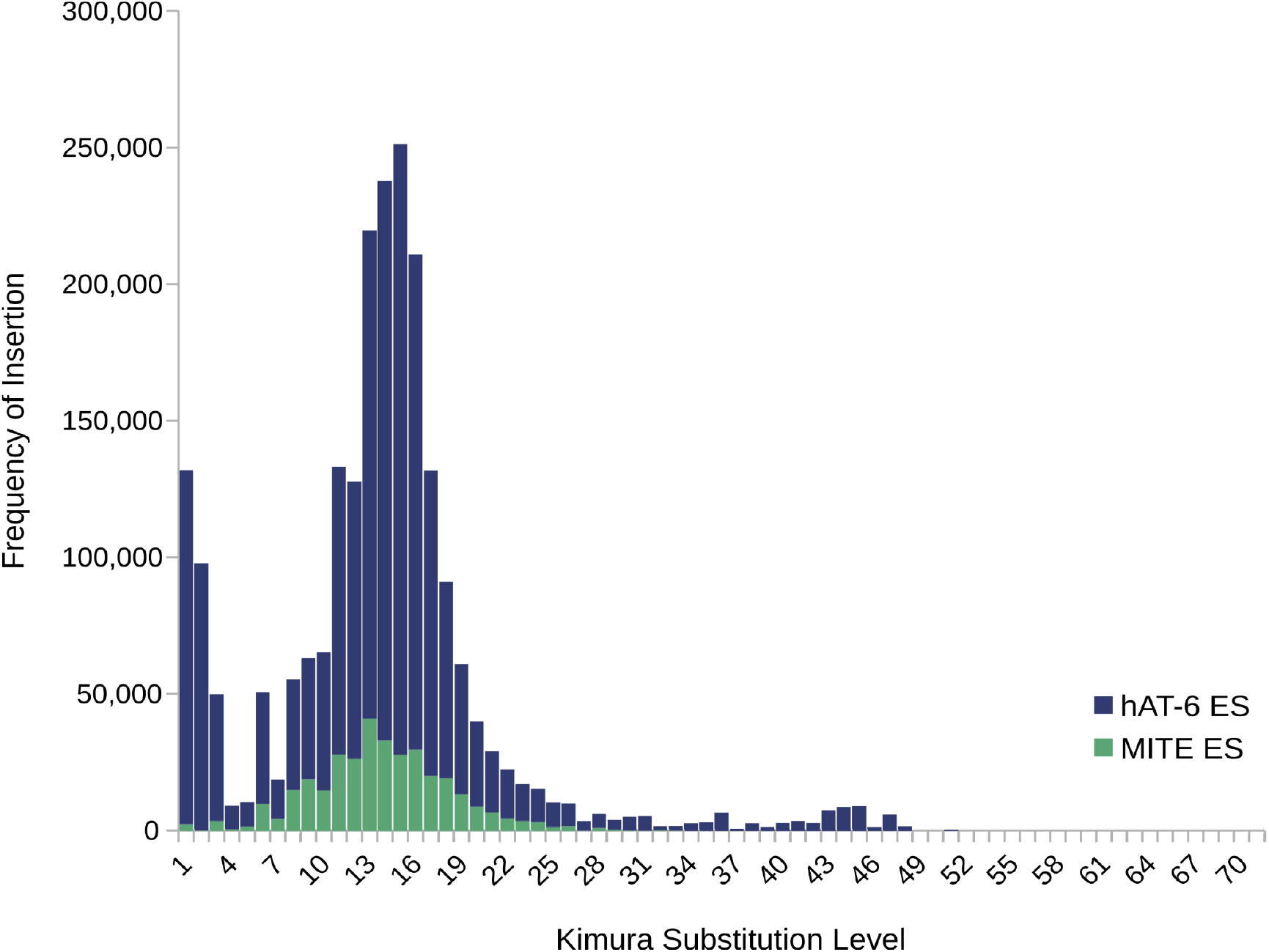
Kimura distance-based divergence of hAT-6 derived MITE compared to hAT-6 from the genome of *Etheostoma spectabile*. The left-hand axis indicates the frequency of TE insertion into the genome and the bottom axis displays relative age shown from left to right.

In Figure 6, we show the frequency of hAT-6 insertion and divergence relative to other vertically inherited DNA TEs and retrotransposons in turtles. In both *M. t. terrapin* and *T. s. elegans*, hAT-6 is present in low quantities compared to other elements but shows less than 1% divergence, further supporting the case for HTT. When comparing hAT-6 to hAT-3, a DNA TE shared by both *M. t. terrapin* and *T. s. elegans*, we see that the hAT-6 Kimura divergence and abundance are more similar than expected between the two turtles, despite the divergence between the two species estimated to be 14.5-15.6 MYA [26]. In addition, hAT-3s from both turtles are nearly identical in sequence but show a higher kimura divergence over time which is consistent with an ancestral repeat. We also show that DNA TEs and retrotransposable elements have similar patterns of copy number increase and divergence, likely indicating the presence of population bottlenecks leading to fixation of insertions [27]. It is important to note that other vertically inherited TEs show peaks consistent with reactivation over time in the turtle genomes, which is not observed with hAT-6 even though hAT-6s in turtles appear to have two peaks (fig. 6). When investigating the sequences corresponding to the second peak between 34%-52% divergence for hAT-6, manual curation revealed that the alignments showed no resemblance to hAT-6, but did resemble other hATs. We suggest they are remnants of other degraded and ancient hATs. Taken together, our results show HTT between distant species but also how horizontal transfer looks in terms of divergence times in turtles and how it corresponds to what is known about their genome evolution.

**Figure 6:**
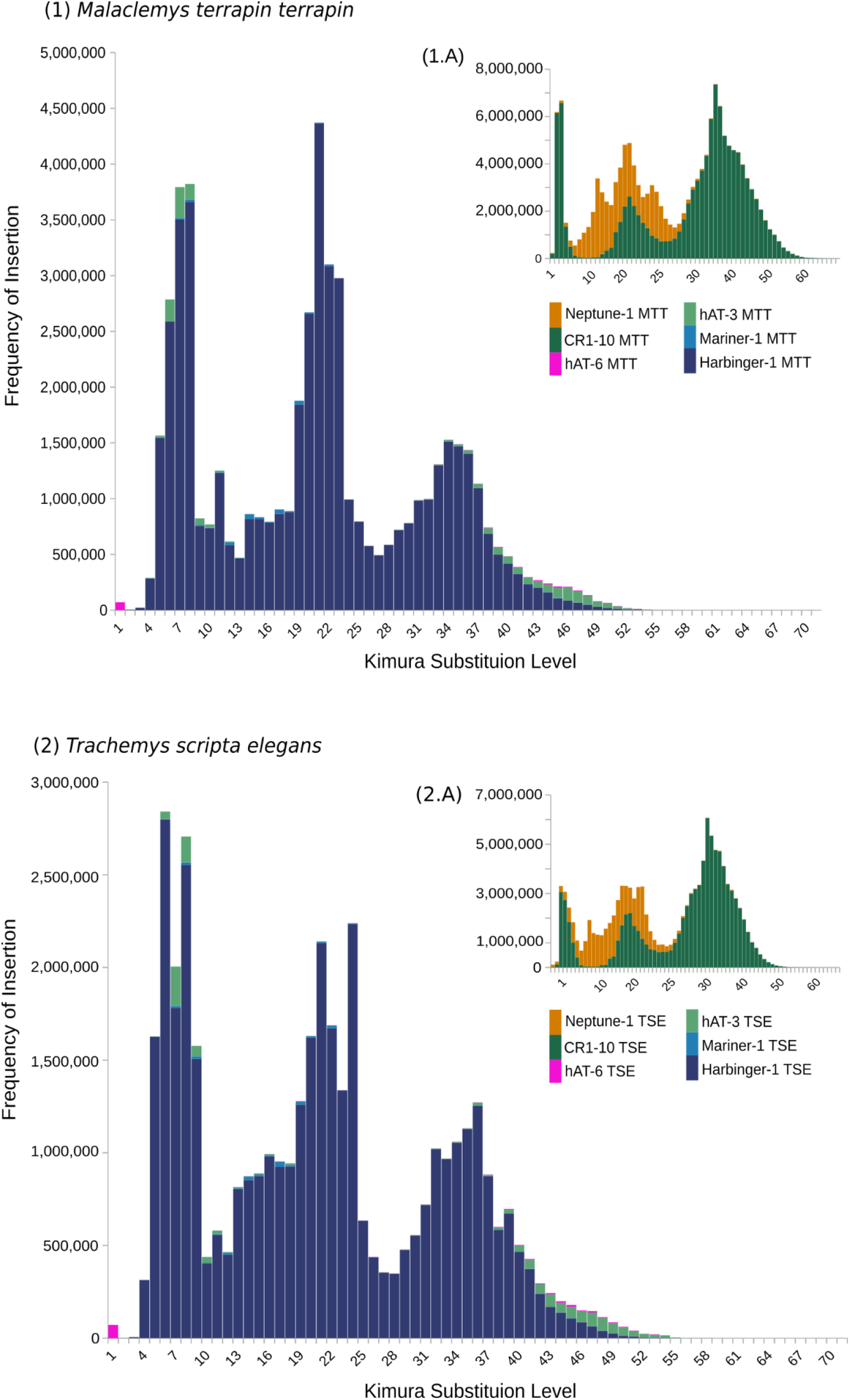
Kimura distance-based divergence of DNA transposons (left plot) and retrotransposons (top right plot) from (A) *M. t. terrapin* and (B) *T. s. elegans*. DNA transposon (hAT-6, hAT-3, Mariner-1, and Harbinger-1) divergence is shown in contrast to the divergence of retrotransposons (Neptune-1 and CR1-10) for both genomes. The left-hand axis indicates the frequency of TE insertion into the genome and the bottom axis displays relative age shown from left to right.

### Aquatic environments and parasite-host relationships may facilitate HTT

The high percentage identity and low divergence of hAT-6 between species are likely the result of independent transfers from an unknown donor(s). The species in this study are largely found in semi-aquatic or aquatic environments but are geographically distant. This indicates possible donor(s) that is/are likely ubiquitously aquatic and/or parasitic in nature. This finding also supports the frequent and recent transfer of transposons in aquatic environments [18,19]. Horizontal transfer is known to be facilitated by parasite-host relationships with recent findings showing HTT of a retrotransposon from parasitic nematodes [21,28]. Pathogens affecting fish, frogs and turtles include nematodes, tapeworms, flukes, trematodes, and myxosporean parasites [22–24]. The myxozoan endoparasite of the common carp (*T. kitaueI*) contained a hAT DNA transposon with motifs remarkably similar to the horizontally transferred hAT-6 TSDs, TIRs, and identical DDE/RW residues. Although at present, there is insufficient evidence to explain the HTT of hAT-6 via a parasitic intermediate, *T. kitauel* shows potential as a parasitic donor as it is known to infect fish and possibly frogs and turtles. In addition to myxozoan parasites as a possible vector, there is a geographical overlap between the turtles and *E. spectabile* (Orangethroat darter) suggesting hAT-6 HTT into turtles may have occurred from *E. spectabile* into turtles through a shared parasitic vector, such as darter fish parasites as all three species have overlapping geographical distributions in North America [29]. But as there is no geographical overlap between *X. tropicalis* and the other fishes, more genomic data from geographically/ecologically overlapping species are required to determine possible donors (Supplementary materials).

## Conclusions

Overall our study expands our knowledge of HTT in aquatic species and especially the evolution of HTT repeats in the slow-evolving genome of turtles. We have documented a rare horizontal transfer event between turtles, ray-finned fishes, and a frog, showing that HTT may be more common than expected in turtles. Our findings support the notion that HTT is a common occurrence in ray-finned fishes and that aquatic environments may facilitate a large number of HTT events. The direction of HTT of hAT-6 in turtles is likely from *E. spectabile* as the species share habitat and overlap geographically. While the donors for HTT are unknown, our results and others from the literature suggest the existence of a cryptic aquatic network of horizontal transfer that is widely distributed given the geographic distances between the HTT recipients.

## Methods

### Identification and classification of horizontally transferred DNA transposon candidates

We performed an *ab initio* repeat annotation for the genome of *M. t. terrapin* using the Comprehensive *ab initio* Repeat Pipeline (CARP) [30]. We identified one DNA TE as a horizontal candidate in *M. t. terrapin* which was originally curated in *X. tropicalis* (Western clawed frog) as hAT-6 XT. We thus renamed the DNA TE in *M. t. terrapin* as hAT-6 MTT. To find more similar sequences to hAT-6 MTT, we performed an extensive local alignment using hAT-6 MTT as a query against the following taxonomic groups: crocodilians, birds, frogs and toads, snakes, turtles, and fish using sensitive BLASTN 2.7.1+ parameters [31] (word-size: 7, match/mismatch score: 4, -5) against RefSeq Representative Genomes [32]. A cutoff of 75% identity and 90% coverage to hAT-6 MTT was used to find full-length transposon sequences (-blastn -e-value 1e-10). We did global alignments of each sequence back to the query species using MAFFT v7.450 using NCBI coordinates to detect TIRs/TSDs [32,33]. We classified sequences as full-length transposons or fragments based on the presence of TIRs and TSDs. Through this process, 8 sequences similar to hAT-6 MTT were found. Open reading frames (ORFs) were identified using GENSCAN and a pairwise alignment matrix (PIM) for all 8 sequences was made using BLASTN [34].

### Construction of Repeat Phylogenies

We aligned horizontally transferred hAT-6 repeats and a sample of hAT annotated sequences from RepBase using MAFFT v7.450 [35]; [32,33] (FFT-NS-1 model). We processed the multiple alignments using Clipkit [36] to select conserved regions allowing smaller final blocks, gap positions within the final blocks and less strict flanking positions. Two independent tree-building tools were used: Fasttree 2.1 (General time-reversible (GTR) model) and IQTree (Stamatakis, 2014) (GTR; 1000 bootstraps) [37,38]. All tree files were visualized and edited using iTOL [39].

### Construction of Species Phylogeny

We used TimeTree to construct a phylogenetic tree of a selection of species to represent amphibians, bony fishes, and reptiles. The tree was visualised on the Interactive Tree of Life (iTOL) [39,40]. In the case where a species of interest was not available on TimeTree, we substituted a species from the same clade.

### Protein Structural Analysis of hAT-6s

To determine the protein structure of horizontally transferred hAT-6s, we used an online protein modelling server, I-TASSER (Iterative Threading ASSEmbly Refinement). We determined the structural features of three hAT-6 DNA transposons with a protein sequence exceeding 600 amino acids: hAT-6 TSE (*T. s. elegans*), hAT-6 SF (*S. foromus*), and hAT-6 SA (*S. acus*) [41]. The run times for each query were respectively 60 hr, 60 hr, and 72 hr. I-TASSER output was visualized using PYMOL (The PyMOL Molecular Graphics System, Version 2.0 Schrödinger, LLC). DD/E and RW amino acid residues in transposase sequences were identified using M-COFFEE with alignment to Hermes transposase (Supplementary Materials) [42].

### Divergence and genome coverage of horizontally transferred DNA transposons

To identify orthologous full-length hAT-6 DNA transposon sequences from the genomes they were curated from to ensure reliable annotation, we performed a reciprocal search [18,43]. A custom database was made using the relevant genomes to find the reciprocal best hit using hAT-6 sequences as a query from the genome of interest. TE-Aid (https://github.com/clemgoub/TE-Aid) was used to align each full-length hAT-6 DNA transposon sequence to the genome it was curated from to visualise the frequency of the complete sequence and fragments.

### Kimura distances

We calculated the Kimura 2-parameter distance for all hAT-6 TEs using RepeatMasker version 4.1.5 and the calcDivergenceFromAlign.pl to obtain the relative age distribution of the TEs in the genome (http://www.repeatmasker.org/RMDownload.html). We also calculated the Kimura 2-parameter distance for various repeats for the Testudine genomes (*T. s. elegans* and *M. t. terrapin*) to determine the age of hAT-6 compared to other repeats. We excluded short simple repeat alignments from further analysis.

## Acknowledgments

We would like to thank our lab for their ongoing support and input while putting together this manuscript.

## Declarations

### Ethics approval and consent to participate

Not applicable

### Availability of data and materials

The dataset(s) supporting the conclusions of this article are included within the article (and its additional file(s)).

### Author Contributions

N.T.H, J.D.G, and D.L.A designed research; J.D.G provided scripts; N.T.H and D.L.A wrote the paper with input from J.D.G

