## Supplementary material for "Jumping between Turtles, Fishes, and a Frog: The Unexpected Horizontal Transfer of a DNA Transposon": HTT_Kimura.docx

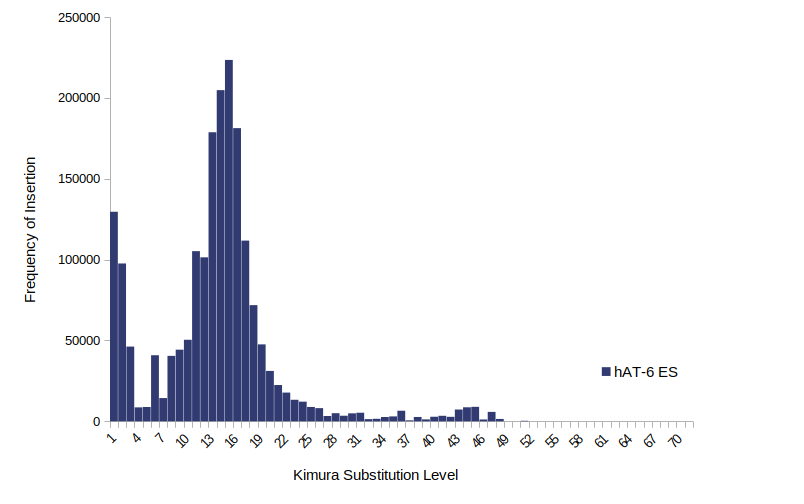

**Figure 1:** Kimura distance-based divergence of hAT-6 from the genome of *Etheostoma spectabile*. The left-hand axis indicates the frequency of TE insertion into the genome and the bottom axis displays relative age with low to high age shown from left to right.

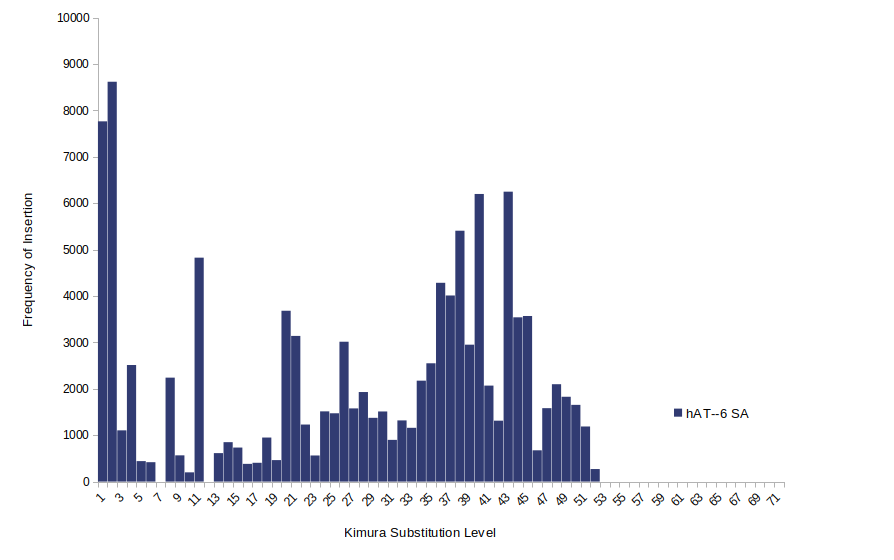

**Figure 2:** Kimura distance-based divergence of hAT-6 from the genome of *Sygnathus acus*. The left-hand axis indicates the frequency of TE insertion into the genome and the bottom axis displays relative age with low to high age shown from left to right.

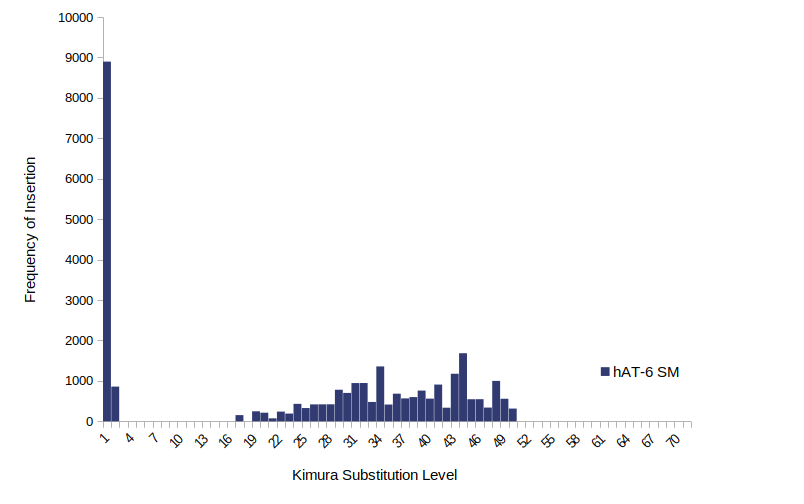

**Figure 3:** Kimura distance-based divergence of hAT-6 from the genome of *Scopthalmus maximus*. The left-hand axis indicates the frequency of TE insertion into the genome and the bottom axis displays relative age with low to high age shown from left to right.

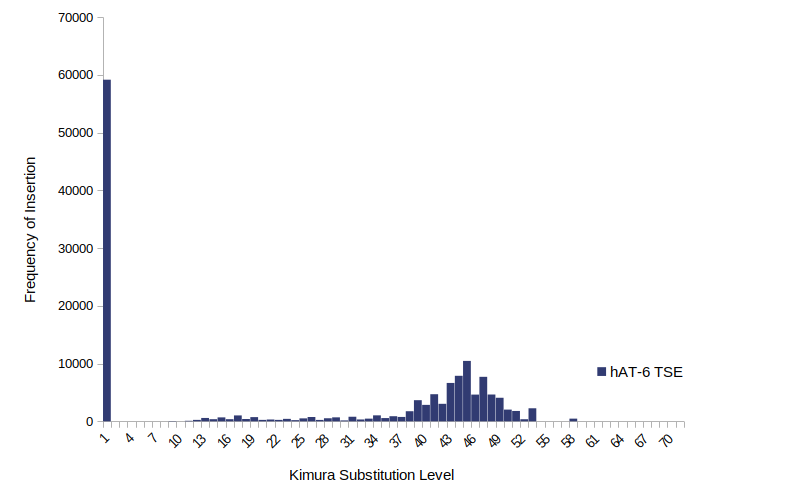

**Figure 4:** Kimura distance-based divergence of hAT-6 from the genome of *Trachemys scripta elegans* The left-hand axis indicates the frequency of TE insertion into the genome and the bottom axis displays relative age with low to high age shown from left to right.

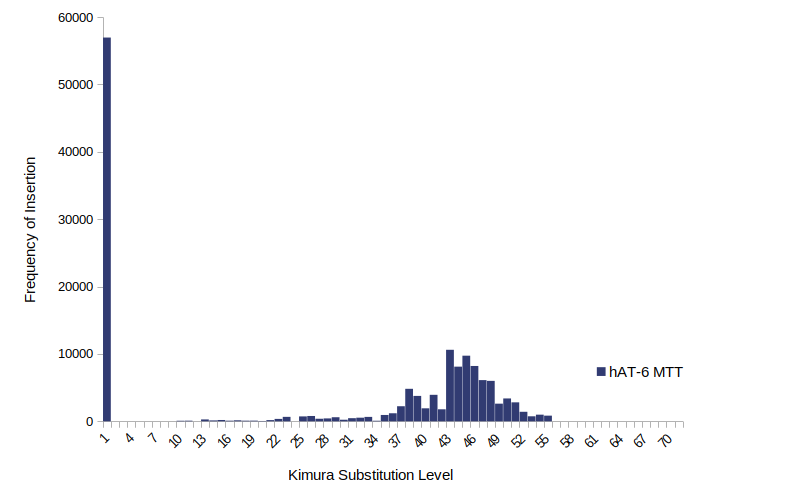

**Figure 5:** Kimura distance-based divergence of hAT-6 from the genome of *Malaclemys terrapin terrapin*. The left-hand axis indicates the frequency of TE insertion into the genome and the bottom axis displays relative age with low to high age shown from left to right.

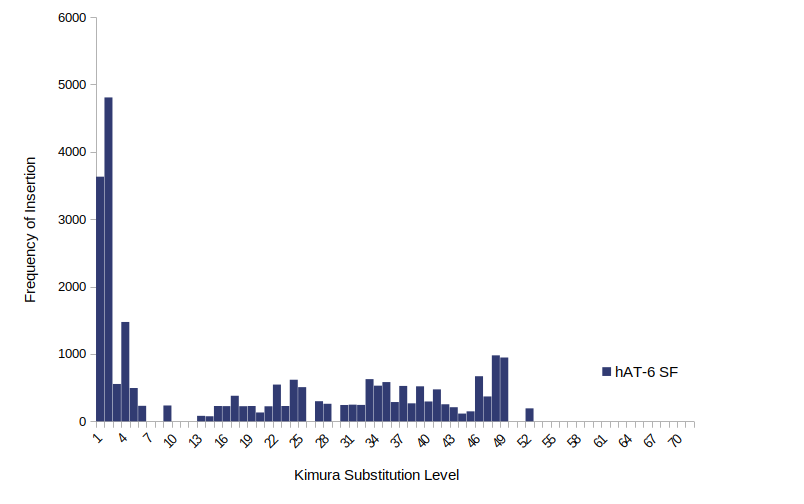

**Figure 6:** Kimura distance-based divergence of hAT-6 from the genome of *Scleropages foromsus*. The left-hand axis indicates the frequency of TE insertion into the genome and the bottom axis displays relative age with low to high age shown from left to right.

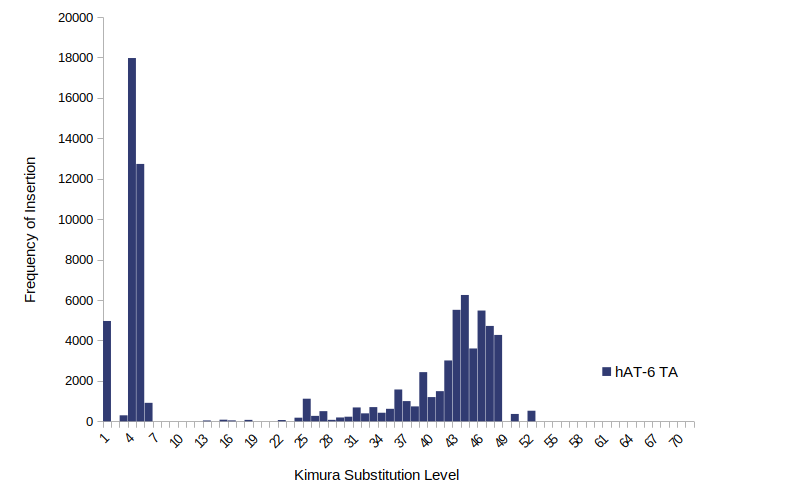

**Figure 7:** Kimura distance-based divergence of hAT-6 from the genome of *Thalassophryne amazonica*. The left-hand axis indicates the frequency of TE insertion into the genome and the bottom axis displays relative age with low to high age shown from left to right.

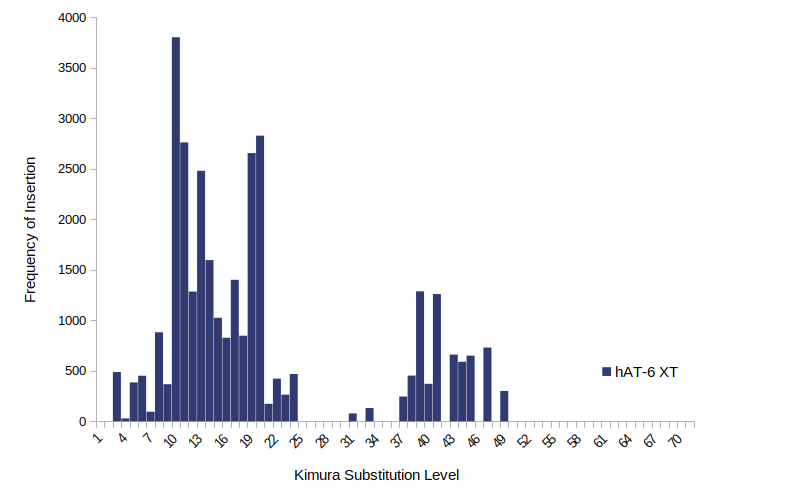

**Figure 8:** Kimura distance-based divergence of hAT-6 from the genome of *Xenopus tropicalis*. The left-hand axis indicates the frequency of TE insertion into the genome and the bottom axis displays relative age with low to high age shown from left to right.
